# The role of working memory for bridging the gap between perception and goal-directed actions: Evidence by mu and beta oscillations in sensorimotor cortex

**DOI:** 10.1101/817742

**Authors:** Daniel Schneider, Marlene Rösner, Laura-Isabelle Klatt, Edmund Wascher

**Affiliations:** Leibniz Research Centre for Working Environment and Human Factors

**Keywords:** Working memory, motor preparation, selective attention, neural oscillations, mu/beta suppression

## Abstract

What mechanisms are at work when transferring a visual representation in working memory into a higher-level code for guiding future actions? We investigated the underlying attentional and motor selection processes in working memory by means of oscillatory EEG parameters. Participants stored two, three or four objects in working memory and subsequent retroactive cues indicated one or two items as task-relevant. The oscillatory response in mu (10-14 Hz) and beta (15-25 Hz) frequencies with an estimated source in sensorimotor cortex contralateral to response side was used as a correlate of motor planning. There was a stronger suppression of oscillatory power when only one item was cued. Importantly, this effect appeared although the required response could not be anticipated at this point in time. This suggests that working memory can store multiple item-specific motor plans and the selection of a stored visual item leads to an automatic updating of associated response alternatives.

## 1. Introduction

In everyday life, we are frequently required to respond to stimuli that are present in our environment. However, what feels natural and automatic, requires a complex cascade of cognitive processes, ranging from sensory perception via motor planning to motor execution. Working memory can be defined as a cognitive stage for the interface between perceptual information, higher-level cognitive operations and goal-directed actions. Based on a retroactive cuing working memory paradigm, this study investigates the nature of action-oriented working memory representation that are required to bridge the gap from a sensory representation to goal-directed actions.

Traditionally, perception and action have been considered to occur in a capacity-limited, strictly-serial fashion^1^. However, this view fails to consider the goal-directedness of the majority of human perception and behavior in every-day life^2^. The theory of event coding proposes a joint mental representation of perceptual information and action plans^2,3^. This means that stimuli and response features are integrated in a way that the presentation of a stimulus leads to perceptual processing as well as to the activation of related response alternatives. Importantly, Hommel, Müsseler, Aschersleben and Prinz^2^ highlighted that a motor plan is a code of the perceptual consequences of a motor action rather than a conscious plan of a complex sequence of muscle tension and relaxation. Important for this account, working memory can not only actively store stimulus representations in different sensory stores^4,5^, but also higher-level representations of to-be-executed actions^6-9^. It is therefore reasonable to assume that the transition from purely sensory information to higher-level sensorimotor representations for guiding future actions proceeds in working memory.

The question how sensory information is used for guiding future actions can be approached by investigating different representational states in working memory as a function of the focus of attention: For example, Cowan postulated that up to four object representations can be held activated in working memory and the focus of attention can be internally directed toward the representation(s) required for an upcoming operation^10^. While working memory contents within the focus of attention are thus stored in a prioritized representational state, irrelevant contents outside the focus of attention are deprioritized and potentially subject to decay and interference by new sensory inputs^11-15^. However, it has so far not been adequately addressed what actually constitutes a mental representation in working memory once it has been selected to guide a future operation. Here, we propose that whenever an upcoming operation is a goal-directed action, working memory representations within the focus of attention can be transferred into a higher-level sensorimotor code for guiding this very action.

An important investigation on the sequence of such sensory- and action-related working memory processes was recently conducted by van Ede, Chekroud, Stokes and Nobre^16^. The authors independently lateralized relevant visual items in a memory array (left vs. right target) and the later required motor responses (button press with left vs. right index finger). Hemispheric asymmetries in alpha oscillations related to attentional selection^17-19^ and in mu/beta oscillations related to planning a left-vs. right-handed response^20-22^ could thus be fully separated from one another. It was shown that these types of oscillatory effects were related to spatially distinct areas of the brain, but occurred in the same time frame. This suggests that attentional selection of visual working memory representations and motor selection or planning, in fact, take place in parallel.

Still, these results could not fully answer the question what actually constitutes a higher-level (motor) representation in working memory. This is because the sensorimotor mu/beta asymmetries prior to already defined motor actions might either reflect a subjective state of the readiness to respond, or the evolving motor plan. First results in support of the second assumption came from an earlier study by Schneider, Barth and Wascher^23^. Contralateral mu/beta suppression as a correlate of the preparation of a working-memory-guided action prior to the presentation of a memory probe was evident when the to-be-reported target feature, but not the required motor action itself was yet defined. In the current investigation, we aimed at further distinguishing the nature of these sensorimotor representations by uncovering the neurocognitive mechanisms underlying the transition from a sensory to a higher-level (goal oriented) mental representation in working memory.

This was done by means of a retroactive cuing working memory task that included spatial cues for indicating either one or two visual objects (discs of different color) as task-relevant after encoding. As against the described prior investigations^16,23^, only the memory probes defined how to further proceed with the cued information. Participants had to indicate whether the memory probe did or did not match in color to (one of) the previously cued item(s) (see figure 1). We further measured EEG correlates of action planning by means of the suppression of mu (10-14 Hz) and beta (15-25 Hz) oscillatory power over sensorimotor cortex contralateral to response side^20-22^ (here: right-handed button presses with the index or middle finger). We hypothesized that the evolving of a motor plan should be evident simultaneous to EEG correlates of attentional selection following the retro-cues that should be reflected in posterior modulations of alpha power (8-14 Hz)^24-28^. The most important question, however, concerns the nature of such motor plans when the probe information and the to-be-executed actions are yet unknown. We propose that working memory is able to store multiple item-specific motor plans or item-response mappings. For example, when cued for selectively storing the color red, the resulting response mapping will contain the following information: ‘If the memory probe is red, press the left button. If the memory probe is not red, press the right button’. These item-response mappings will vary in complexity as a function of the number of working memory items retroactively cued. As the suppression of contralateral mu/beta power reflects the efficiency in preparing for an upcoming action^22,29^, it should be strongest for unambiguous one-item response mappings. These results would indicate that working memory manages the transfer of sensory inputs into sensorimotor representations in order to prepare for the goal-directed behavioral interaction with a complex and usually not entirely predictable environment.

**Figure 1.**
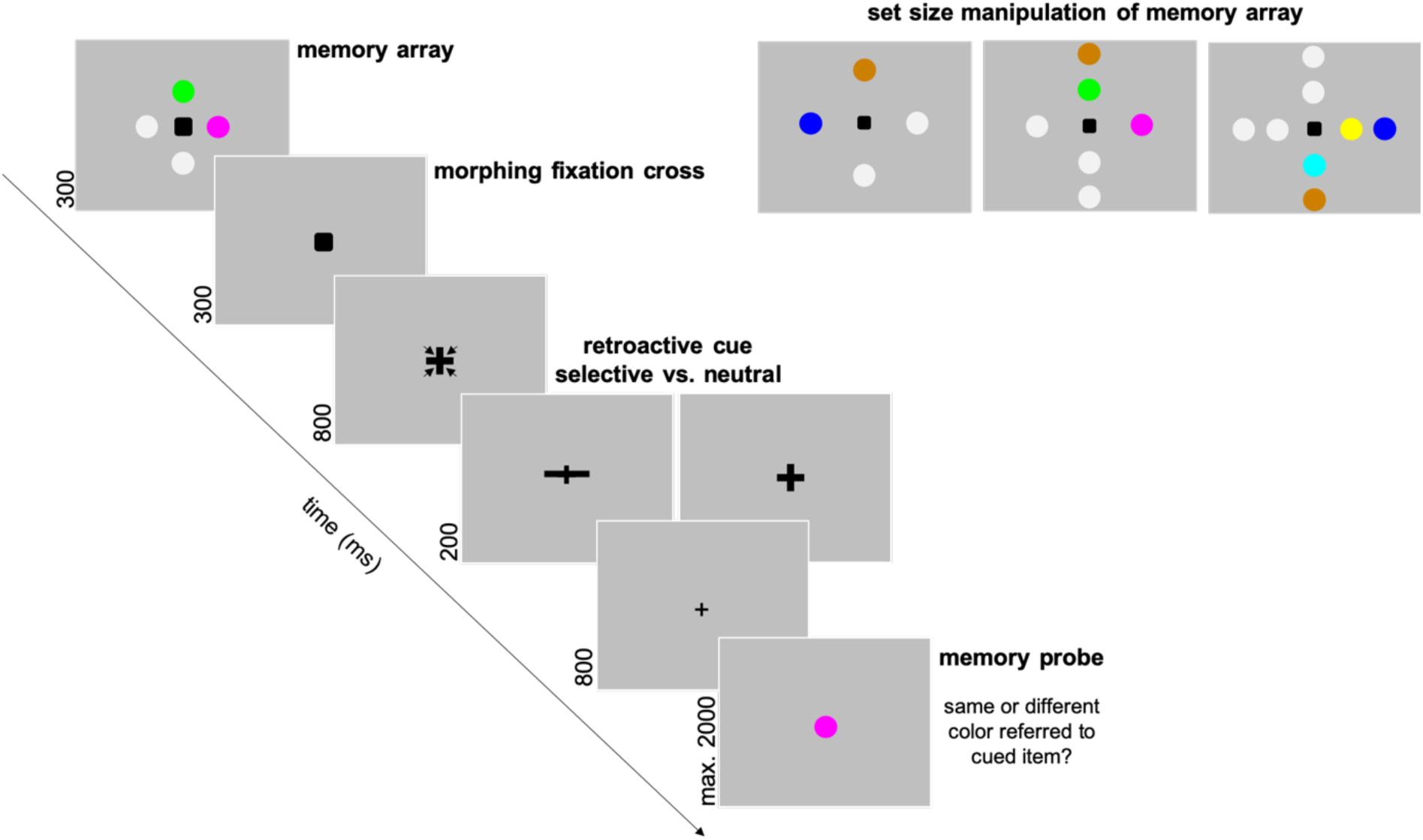
Experimental design. Participants had to store a varying number of colored discs in working memory and were subsequently cued to remember only a subset of the colors presented. When the central memory probe was presented, participants had to indicate whether its color matched or did not match the color of the cued item(s). Either cued or non-cued items were presented on the lateral positions.

## 2. Results

### 2.1 Behavioral results

Twenty-four healthy participants performed in a working memory experiment including retro-cues that indicated the relevance of already stored items for a later recognition task. Participants were first required to remember differing numbers of colored disks presented in a memory array (set size: two, three or four items) and to shift attention to a subset of the initially stored information based on the retro-cues. Based on this experimental design, the number of cued items (one vs. two) and the number of non-cued items following the retro-cues (one vs. two) varied independently. Furthermore, we implemented a condition with a neutral retro-cue that indicated all items of a two-item memory array as further on task relevant. Subsequently, participants indicated by button press whether the color of a centrally presented memory probe did or did not match the cued item (one-item cued condition) or one of the cued items (two-items cued condition). The type of response required was thus not predictable prior to presentation of the memory probes.

Lower error rates were shown when only one item was cued, *F*(1,23)=48.169, *p*<0.001, *p*_*crit*_=0.033, *η*^*2*^*p*=0.677, and for one compared to two non-cued mental representations, *F*(1,23)=48.701, *p*<0.001, *p*_*crit*_=0.05, *η*^*2*^*p*=0.679. Additionally, the effect of the number of non-cued items was further modulated by the number of cued items, *F*(1,23)=24.367, *p*<0.001, *p*_*crit*_=0.017, *η*^*2*^*p*=0.514. Post-hoc analyses indicated that the error rate difference between the conditions with one compared to two non-cued items was reduced when only one, *t*(23)= −3.250, *p*_*adj*_=0.004, *d*_*z*_=−0.663, compared to two working memory items was indicated as relevant, *t*(23)=−7.804, *p*_*adj*_<0.001, *d*_*z*_=−1.593 (see figure 2).

**Figure 2.**
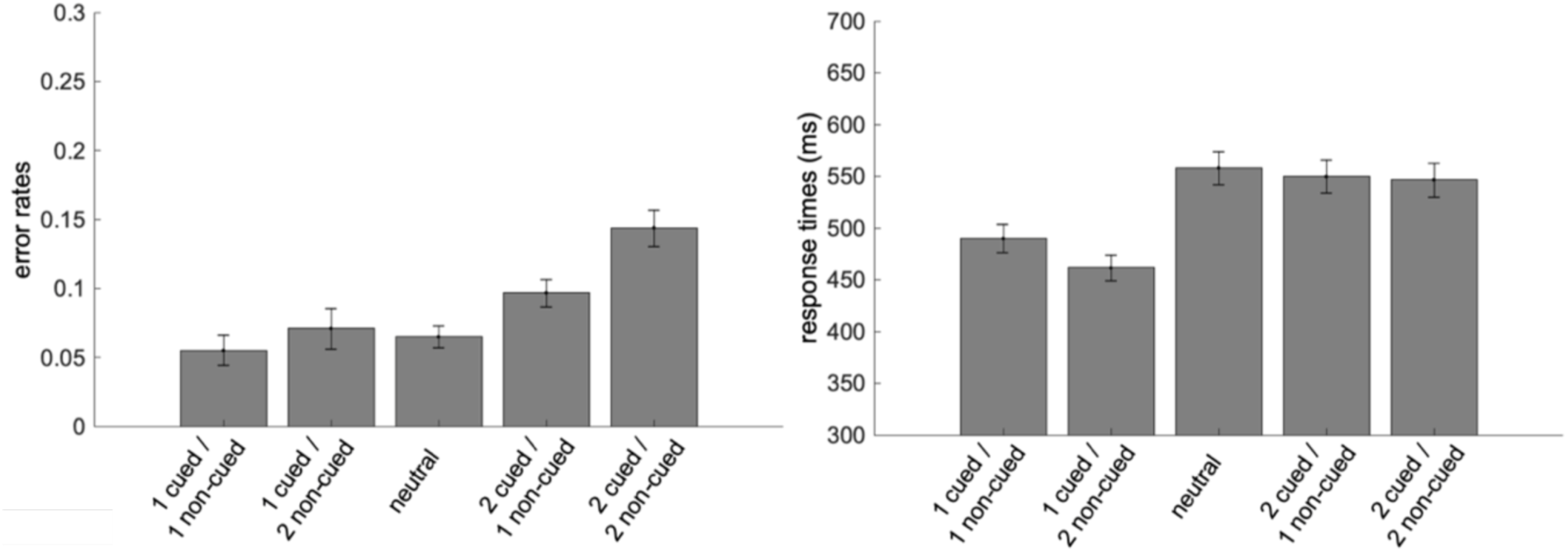
Error rates and response times depending on retro-cue positions. The error bars reflect the standard error of the mean.

Also RTs revealed a significant interaction of the number of cued and non-cued items, *F*(1,23)=20.014, *p*<0.001, *p*_*crit*_=0.033, *η*^*2*^*p*=0.465, as well as significant main effects for these factors. Faster RTs were evident for one compared to two cued items, *F*(1,23)=97.035, *p*<0.001, *p*_*crit*_=0.05, *η*^*2*^*p*=0.808, and for two compared to one non-cued item, *F*(1,23)=19.604, *p*<0.001, *p*_*crit*_=0.017, *η*^*2*^*p*=0.460. The interaction was based on the fact that the effect of the number of non-cued items was only evident when one item was retroactively cued, *t*(23)=5.044, *p*_*adj*_<0.001, *d*_*z*_=1.030 (two-items cued: *t*(23)=0.983, *p*_*adj*_=0.336, *d*_*z*_=0.201).

Furthermore, performance in the experimental condition with a two-item memory array was analyzed as a function of retro-cue type (selective vs. neutral cue). A benefit for the selective retro-cues was found on the level of RTs, *t*(23)=−7.149, *p*<0.001, *d*_*z*_=−1.459, but not for error rates, *t*(23)=−0.893, *p*=0.381, *d*_*z*_=−0.182.

### 2.2 EEG results

#### 2.2.1 EEG correlates of attentional selection in working memory

In the first place, we analyzed the oscillatory response following the memory arrays dependent on the lateral positions indicated as relevant or irrelevant by means of the retro-cues. These analyses included conditions with lateral target items and central non-cued items, as well as conditions with non-cued information at the lateral positions and relevant items at central positions. These conditions were further divided into conditions with one vs. two cued items and one vs. two non-cued items. Additionally, there was a neutral retro-cue condition with two memory array items that were both cued as further on task-relevant. The orienting of attention following the retro-cues was assessed by measuring the contralateral and ipsilateral portions of the oscillatory EEG signal in alpha frequency range (8 – 14 Hz) referred to the left and right cued vs. non-cued positions (for further details, see methods section). Importantly, these effects were not confounded by EEG correlates of motor planning, as stimulus positions and the side of responses were independent.

As indicated in figure 3, contralateral minus ipsilateral event-related spectral perturbations (ERSP; PO7/8, P7/8, PO3/4, P5/6) in the alpha frequency range (8-14 Hz average) revealed hemispheric asymmetries already following memory array presentation. These effects should be related to the orienting of attention toward the positions of the task-relevant lateral items and a subsequent re-orienting of attention in anticipation of the central retro-cues. Importantly, while these asymmetric patterns did not differ between cued vs. non-cued lateral items, the alpha power asymmetries following retro-cue presentation revealed opposing effects for these experimental conditions. This difference was statistically reliable within a time range from 1876 to 2198 ms (i.e., 476 – 798 ms after retro-cue onset). A post-hoc analysis of variance (ANOVA) with the factor ‘eliciting stimulus’ (cued vs. non-cued item(s) lateral) and the number of cued items and non-cued items as within-subject factors did not reveal any significant interactions (all *p*-values > 0.1). Furthermore, only the increase in alpha power contralateral to the positions of non-cued items differed significantly from the hemispheric alpha power asymmetry in the neutral condition (see figure 3). This effect appeared in an interval from 1856 to 2310 ms (i.e., 456 – 910 ms after retro-cue presentation).

**Figure 3.**
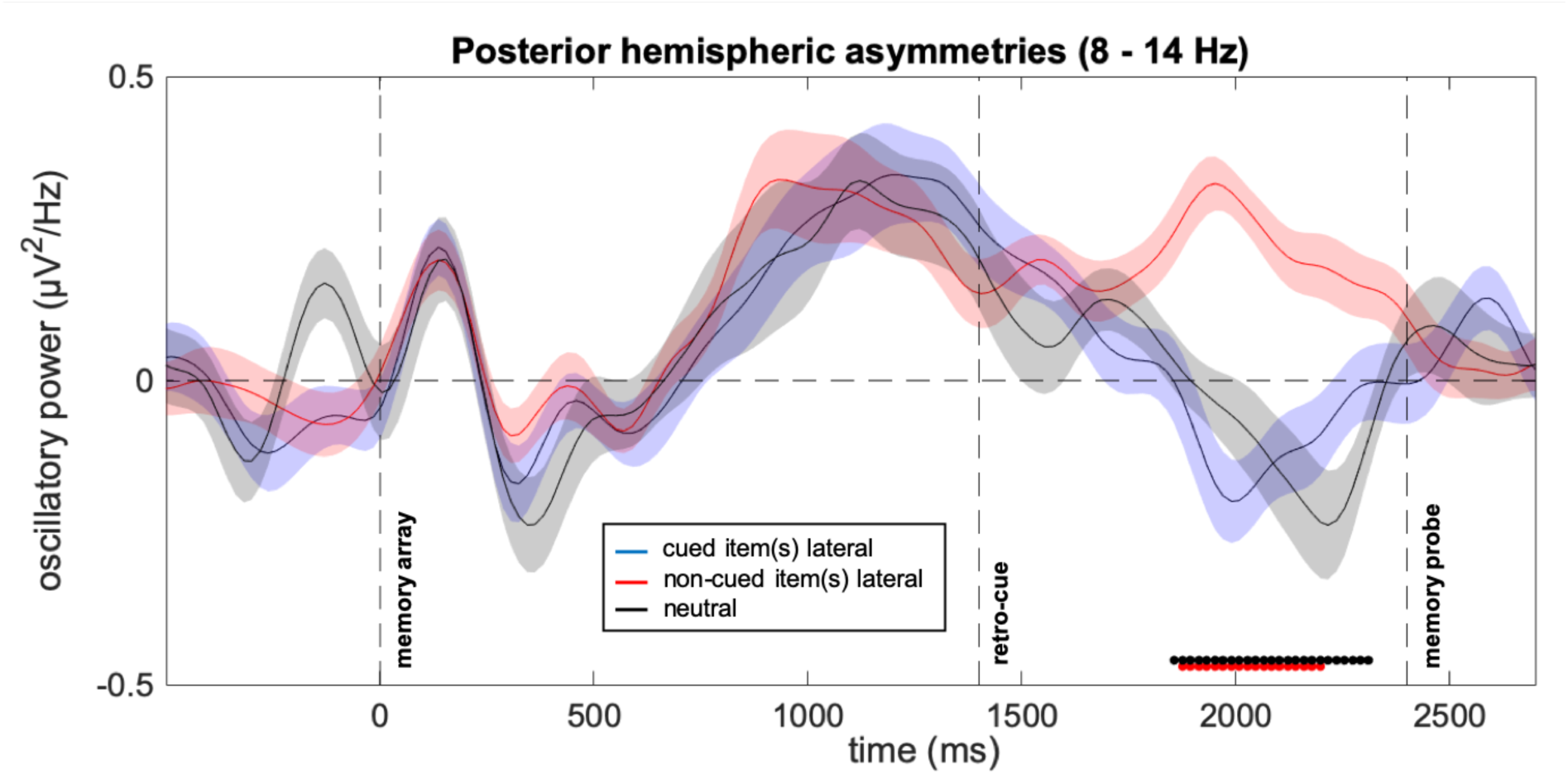
Hemispheric alpha power asymmetries dependent on the lateral position of the memory items. The contralateral minus ipsilateral differences in posterior alpha power (8-14 Hz) is displayed for conditions with cued vs. non-cued lateral items and for the neutral retro-cue condition. The black and red dotted lines reflect significant clusters when comparing conditions with lateral non-cued items to those with lateral cued items (red) and neutral retro-cues (black). The shaded areas around the plotted curves reflect the standard error of the mean.

#### 2.2.2. EEG correlates of motor planning

In order to isolate the modulation of mu and beta oscillatory power in preparation for upcoming actions from modulations of posterior alpha power following retro-cue presentation, the clustering of independent components (ICs) within the EEG signal^30,31^ has proven to be a powerful tool^9,23^. We clustered ICs on the basis of scalp topographies, dipole estimates, ERSPs and frequency spectra (see methods section for more details). Two clear clusters for the left-hemispheric (29 ICs from 19 participants) and right-hemispheric mu/beta oscillatory response (24 ICs from 17 participants) were generated. Dipole density analysis, as included in the EEGLAB toolbox for MATLAB, indicated likely neural sources in left- and right-hemispheric postcentral and precentral gyri (see figures 4 and 5).

**Figure 4.**
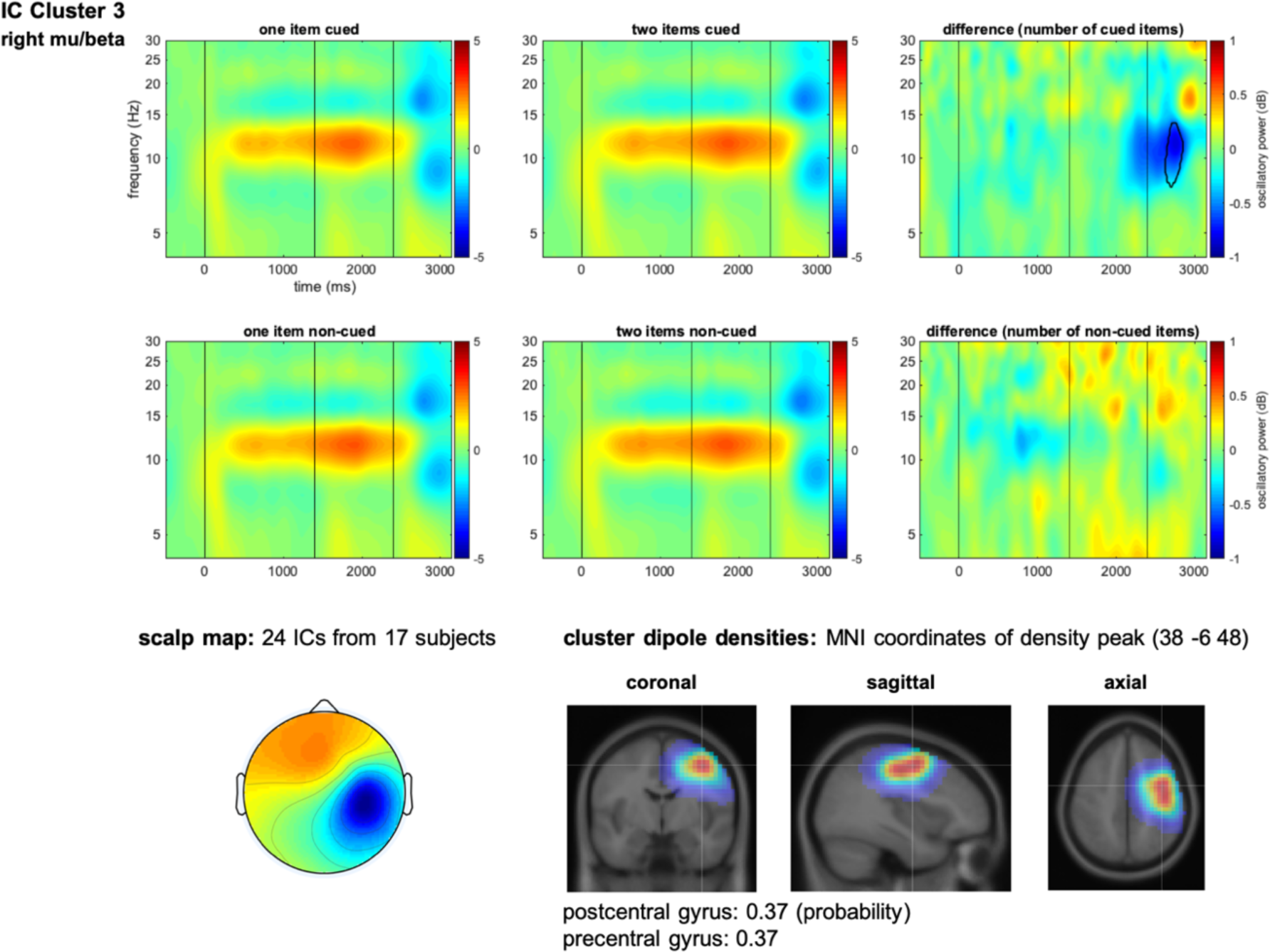
Event-related spectral perturbations (ERSPs) for the right-hemispheric mu/beta cluster. The upper ERSP plots show the ERSPs as a function of the number of cued items (one vs. two). The lower plots show the ERSPs averaged across conditions with one vs. two non-cued items. Statistically significant clusters are highlighted in the difference plots. The lower part of the figure shows the average scalp map and dipole density plots of the IC cluster. The vertical lines indicated the time points of memory array, retro-cue and probe display onsets.

**Figure 5.**
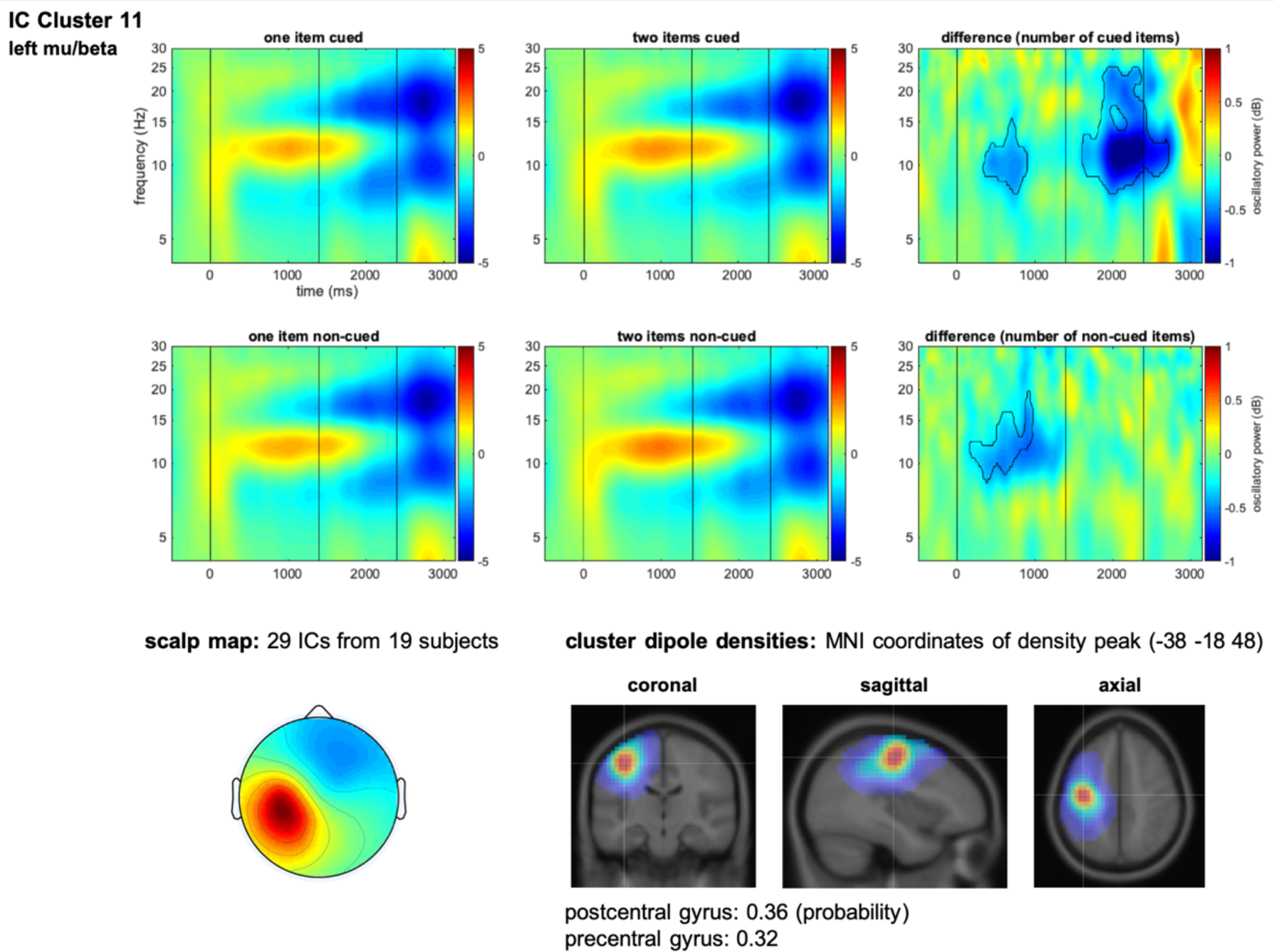
Event-related spectral perturbations (ERSPs) for the left-hemispheric mu/beta cluster. The upper ERSP plots show the ERSPs averaged across all conditions with one-item retro-cues and cues indicating two-items as further on task relevant. The lower plots show the ERSPs as a function of the number of non-cued items (one vs. two). Statistically significant clusters are highlighted in the difference plots. The lower part of the figure shows the average scalp map and dipole density plots of the IC cluster. The vertical lines indicated the time points of memory array, retro-cue and probe display onsets.

The ERSPs of these IC clusters were compared as a function of the number of cued and non-cued items following the retro-cues. For both clusters, there was an increase in oscillatory power from ∼10-14 Hz that started shortly after presentation of the memory arrays. In the right-hemispheric mu/beta cluster, this effect was evident independent from the number of cued or non-cued items. An increased suppression of oscillatory power in the mu frequency range was evident only after memory probe presentation (see figure 4). In the left-hemispheric cluster, alpha power increased with both the number of cued and non-cued items already prior to retro-cue presentation, suggesting a relation to the varying set-sizes of the memory array. More important for the current purposes, there was a suppression of oscillatory power with a peak in both higher alpha and beta frequencies for the left-hemispheric cluster that appeared after retro-cue, but clearly before memory probe presentation (i.e., a typical characteristic for oscillatory correlates of motor preparation and imagery^20,32,33^). This effect was exclusively modulated by the number of cued items, with a stronger suppression of oscillatory power prior to memory probe onset for the one-item cued condition. This result pattern highlights that, although decisional processes could only be based on the memory probes, the presentation of the retro-cues led to an early refinement of motor representations in anticipation of the to-be-given responses.

When comparing the time courses of the attentional modulations on posterior alpha power asymmetries (see figure 3) and the suppression of contralateral mu/beta power as a function of the number of working memory items cued (see figure 5), it can further be concluded that the underlying processes proceeded in parallel. In an analysis with average mu/beta power from 10 – 25 Hz, a reliable effect of the number of cued items was observed in an interval from 1800 to 2538 ms (i.e., 400 – 1138 ms after retro-cue onset).

#### 2.2.3. Brain-behavior correlations

If the suppression in oscillatory power prior to memory probe presentation truly reflects motor preparation, there should be a relation between this effect and the speed of responses to the memory probes. Participants with fastest RTs should thus also feature a stronger preparatory suppression of mu/beta oscillatory power in the sensorimotor cortex contralateral to the side of the to-be-given responses. This was approached by correlating the oscillatory response in the left-hemispheric mu/beta IC cluster averaged across experimental conditions with the average individual RTs (for correct responses only). Correlation coefficients were calculated for all time and frequency combinations and statistical significance was again assessed by means of a cluster-corrected permutation approach (for further details, see methods section). As indicated in figure 6, there were two clear clusters with positive between-subjects correlations between the amplitude of mu and beta power suppression and subsequent RTs.

**Figure 6.**
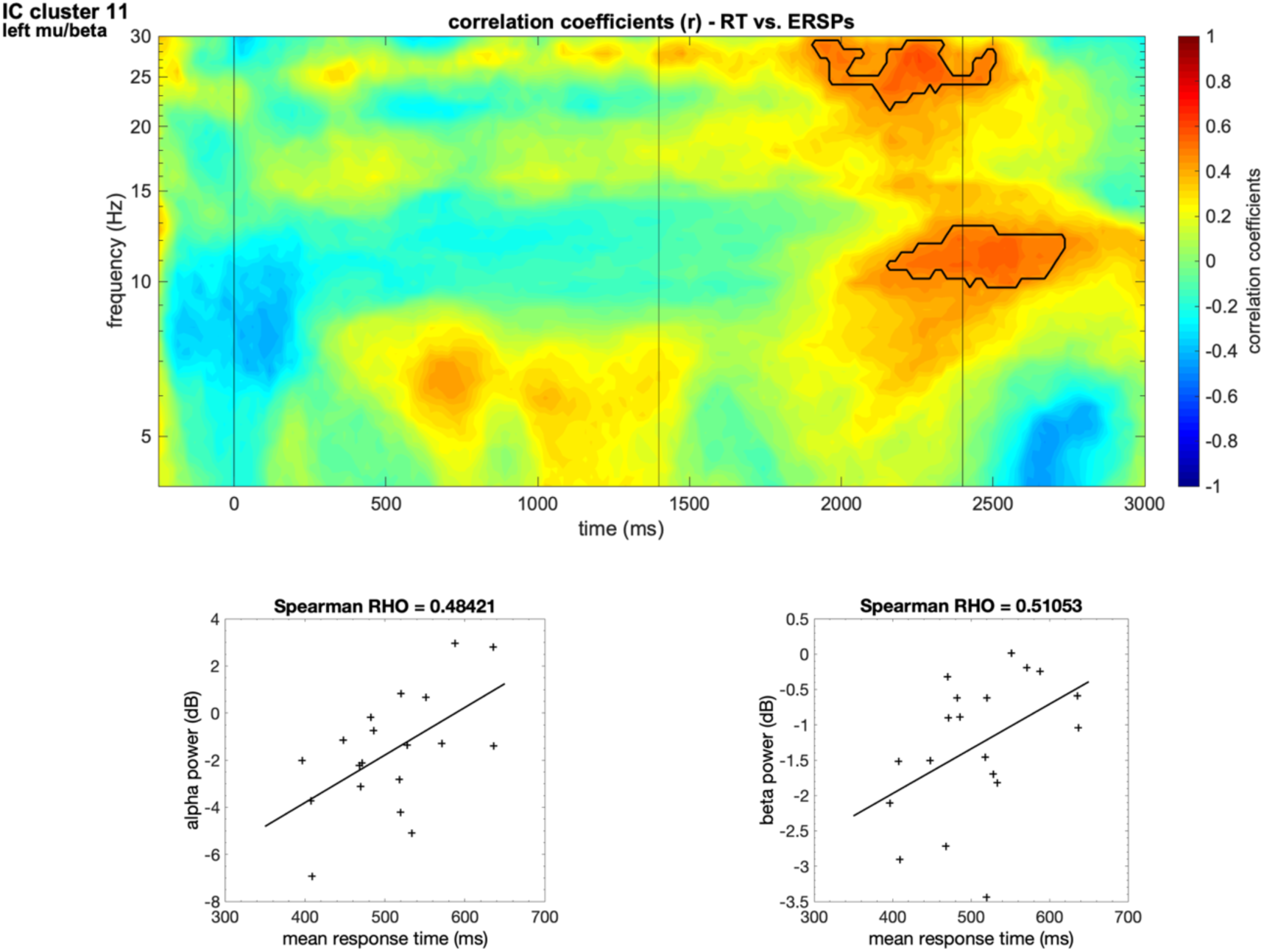
Between-subjects correlation of oscillatory power and average response times. Two clusters with reliable correlations were evident, one in mu frequencies (∼10-13 Hz) and one in higher beta frequencies (∼23-30 Hz). Both correlations appeared prior to memory probe presentation. The vertical lines indicated the time points of memory array, retro-cue and probe display onsets. The lower part of the figure shows the correlations averaged across the significant clusters.

## 3. Discussion

By means of oscillatory parameters in the EEG, the present study examined the neurocognitive processes underlying the transition from sensory working memory representations to higher-level sensorimotor codes for guiding goal-directed actions. We conducted a retro-cue-based working memory experiment that required to selectively store either one or two colors for a later comparison with a memory probe. Error rates increased as a function of the number of both cued and non-cued items, suggesting a general performance deficit with increasing set-sizes of the memory array. This decrease in performance was reduced when retro-cues indicated only one working memory item as task-relevant, in line with prior investigations showing that retro-cue benefits are strongest with a one-item focus of attention^34,35^. This is further supported by the RT effects, with fastened RTs when only one item was retroactively cued (see figure 2).

The retro-cue benefits found on behavioral level can be explained by the electrophysiological correlates of motor representations emerging following retro-cue presentation. A clustering approach on the level of the ICs of the EEG signal resulted in a left-hemispheric mu/beta cluster contralateral to response side and a right-hemispheric cluster ipsilateral to response side. The estimated neural sources of these clusters were located in the left vs. right sensorimotor cortex. Prior research has indicated that while motor planning processes are predominantly reflected in a contralateral mu/beta suppression, motor execution leads to a suppression of mu/beta power within both hemispheres^36^. Accordingly, the typical double-peak in the suppression of mu/beta power appeared already prior to memory probe presentation for the contralateral IC cluster (see figure 5), but after memory probe onset (during response execution) for the IC cluster ipsilateral to response side (see figure 4).

Mu/beta power contralateral to response side featured a stronger decrease when one item compared to two items was retroactively cued. This effect was not confounded by varying set-sizes of the memory array, as such modulations should have been evident as a function of both the number of cued and non-cued items. Importantly, the stronger decrease of mu/beta power following one-item retro-cues appeared already prior to memory probe presentation and can thus be related to the evolving of a motor representation in working memory. Yet, how could such a motor representation look like based on the current experimental design? Independent of experimental condition, the retro-cues provided no information on how to respond to the memory probes. Only the combination of cued working memory items and the probe stimulus defined the to-be-given responses. For this reason, the readiness to respond or the anticipation of a specific movement that has been shown to be reflected in contralateral mu/beta suppression^22,29^ cannot have differed between the retro-cue conditions prior to memory probe presentations. Furthermore, participants were unable to form a specific representation of their behavioral goal (e.g., ‘rotate a bar until it reaches a certain orientation’) before the memory probes were presented.

This leaves the possibility that mu/beta suppression in the left-hemispheric cluster reflected the activation of multiple response alternatives associated with the stored sensory representations. In other words, participants activated a stimulus-response (S-R) mapping that contained information about both the selected sensory representations and the associated response alternatives in anticipation of the memory probes. The S-R mappings following a one-item retro-cues were less complex than the assignment of response alternatives to two colored items, in line with the above-reported behavioral results showing faster RTs when one item was cued. Mu/beta suppression in contralateral sensorimotor cortex might thus reflect the unambiguity of S-R mappings that are activated to guide future actions. This notion corresponds to the theory that mu oscillations in sensorimotor cortex are related top-down modulations by the frontal mirror neuron system and fulfill an important role for the bridging between perception and action^37,38^. For example, prior research has shown that mu and beta power decreases in sensorimotor and motor cortex not only during action planning and execution, but also when observing an action^39-41^. Thus, the sensorimotor system is optimally suited for representing and potentially linking stimulus information and action plans.

Further support for this notion comes from prior investigations showing that it is possible to simultaneously plan reaching movements towards multiple target objects and to subsequently only select one out of the already planned movements^7,42,43^. Referred to the current experiment, the S-R mapping following a one-item retro-cue toward a red item would contain the following information: ‘If the memory probe is red, press the left button, if not, press the right button’. In this sense, the presentation of the memory probes triggered the selection of an already prepared motor action. Accordingly, participants that were more efficient in simultaneously preparing multiple item-specific motor plans, as reflected in stronger suppression of mu/beta power prior to memory probe presentation (see figure 6), responded faster to the memory probes.

Furthermore, it is important to also consider the time course of the transfer of a sensory working memory representation to a higher-level sensorimotor code. Posterior hemispheric asymmetries relative to the position of lateral cued vs. non-cued items were assessed as correlates of the attentional orienting on the level of sensory working memory representations. There was an increase in alpha power over posterior visual areas contralateral to the side of non-cued working memory content and a suppression of alpha power contralateral to the side of the cued item(s) (see figure 3). According to prior findings, a contralateral suppression should reflect the focusing of attention on the cued information, while an increase in alpha power contralateral to irrelevant information should indicate an inhibitory process for withdrawing attention from non-cued working memory content^44,45^. The time course of the hemispheric asymmetries in posterior alpha power provided information about the time required to update the priorities of sensory representations stored in working memory. The orienting of attention toward cued information and the disengagement of attention from non-cued information proceeded simultaneous to the specification of action plans (see figures 3 and 5). We thus propose that the retroactive orienting of attention in working memory leads to a parallel updating of sensory representations and associated motor plans, as also proposed by van Ede and colleagues^16^.

In what way do these results change our current view of working memory? Investigations on the cognitive mechanisms underlying the updating of working memory contents according to their task-relevance usually focus only on the consequences for the sensory representations stored. For example, it was shown that retroactively cued working memory content is less susceptible to sensory distractions^11,13,34,46,47^, more precisely recalled^48,49^ and can bias the perceptual and attentional analysis of new task-relevant inputs^50-52^. Yet, recent research has suggested that there is more to a prioritized working memory representation than this: Rademaker, Chunharas and Serences^53^ showed that activity patterns in intraparietal sulcus (IPS) represented a higher-level memory code that was transformed away from the initial sensory representations stored in working memory. The observation that attended working memory content is less susceptible to sensory interference can thus be explained by the fact that its mental representation is no longer purely sensory in nature. In a comparable way, we argue that the oscillatory patterns in mu/beta frequencies observed here reflected a higher-level working memory representation that was not only based on sensory features, but also entailed associated response alternatives. In line with the investigation by Rademaker and colleagues^53^, the mu/beta oscillations reflecting the combined sensory and motor representations in working memory were not affected by the presentation of sensory distractors^23^.

In summary, we investigated how visual stimuli stored in working memory are transferred into higher-level mental representations for guiding future actions. We found that retro-cues indicating only one item compared to two items as task-relevant entailed a behavioral benefit in response accuracy and RTs (see figure 2). The EEG results indicated that this behavioral benefit might be explained by taking into account the differences in S-R mappings evolving after retro-cue presentation in these experimental conditions. An IC cluster featuring the oscillatory response in mu/beta frequencies with an estimated source in sensorimotor cortex contralateral to response side showed a stronger suppression prior to memory probe presentation when one item was cued (see figure 4). This suppression, associated with the preparation of a motor response, was observed although the required responses could not be anticipated at this point in time. This suggests that it reflected the activation of alternative response plans associated with the selected sensory representations (i.e., S-R mappings). Accordingly, participants with high efficiency in this mechanism featured faster responses to the memory probes (see figure 6). Furthermore, the motor-related effects in mu/beta frequencies proceeded simultaneous to the posterior hemispheric asymmetries reflecting the retroactive attentional orienting towards cued information and the inhibition of non-cued working memory content (see figure 3). This might suggest that attentional selection of working memory representations leads to an automatic updating of the associated response alternatives. Further research should investigate the relation between capacity limits regarding the storage of motor plans in working memory and contralateral mu/beta suppression. As shown by our current findings, this oscillatory correlate of sensorimotor working memory representations might be a valuable indicator for the individual ability to behave purposefully in a not fully predictable environment.

## 4. Materials and methods

### 4.1 Participants

Twenty-four participants (15 females, mean age = 23.5 ±2.8, range = 19 – 30) were tested. They were compensated with 10 € per hour or course credit for their participation. All participants reported right-handedness, which was further confirmed with a handedness questionnaire. None reported any neurological or psychiatric condition and all had normal or corrected-to-normal vision. Color perception was ensured by means of the Ishihara Test for Color Blindness. Before the beginning of the experiment, all participants gave written informed consent. The experimental procedure was approved by the local ethics committee of the Leibniz Research Centre for Working Environment and Human Factors and is accordance with the Declaration of Helsinki.

### 4.2 Stimuli and procedure

The stimuli were displayed on a 22-inch CRT monitor (100 Hz; 1024 × 768 resolution). A maximum of four colored disks (set-size varied between two, three and four items) were presented as memory array stimuli. Disk colors were randomly drawn without replacement from a pool of eight possible colors (RGB values: red: 255 – 0 – 0; blue: 0 – 0 – 255; green: 0 – 255 – 0; yellow: 255 – 255 – 0; magenta: 255 – 0 – 255; cyan: 0 – 255 – 255; purple: 50 – 0 – 100; orange: 205 – 128 – 0; grey: 128 – 128 – 128; average luminance = 53.6 cd/m^2^). The stimuli were presented on a grey background and light-grey irrelevant filler items (144 – 144 – 144, 53.6 cd/m^2^) were shown on the opposite positions (see figure 1).

Participants had to maintain fixation throughout the trials. One or two memory array items were presented on the vertical midline and horizontal midline, respectively. The stimuli closest to the fixation cross (inner position) were located on a hypothetical circle with a radius of 1.25°, while the stimuli on the outer positions were displayed with a 2.5° distance from fixation. After memory array offset, there was a 300 ms delay before the fixation cross morphed into small dot and then changed to the retro-cue after 800 ms. This ensured that participants maintained fixation before the onset of the retro-cues. The cues were presented for 200 ms and indicated which of the two axes of stimuli (vertical or horizontal) would be relevant for the memory task. We further made use of a neutral retro-cue condition where both the vertically and horizontally presented items remained task relevant (only for two-item memory arrays). Depending on the combination of memory arrays (i.e. two, three or four colored disks) and retro-cues, there were five possible conditions: one item cued/one item non-cued, one item cued/two items non-cued, neutral (both items cued), two items cued/one item non-cued and two items cued/two items non-cued. Additionally, either the cued or non-cued items were presented on the lateral positions (left vs. right of fixation). An equal number of trials from each condition was presented to the participants in random order.

Following retro-cue offset and an additional 800 ms delay period, participants were probed with a single colored disk presented in the center of the screen. They were instructed to indicate by button press if the probe color had been presented on the cued position(s). The index and middle finger of the right hand were used for responding and the assignment of response categories (‘same’ vs. ‘different’ memory probe color referred to the cued item(s)) to response keys was counterbalanced across participants. Half of the probe colors were presented on the cued stimulus axis (i.e., cued probes), while 20% were present in the memory array, but on the non-cued positions. The remaining 30% of trials included memory probes presented in a new color respective the prior memory. This difference between the proportion of non-cued and new probes was due to the fact that there were no non-cued items in the neutral retro-cue condition.

The inter-trial interval varied randomly between 500 and 1000 ms. Each participant completed a total of 1440 trials, which were preceded by 20 practice trials. Trials were arranged in a set of 8 blocks containing 180 trials each. The experimental blocks were interrupted by 2-minute breaks to prevent fatigue in the course of the experiment. The procedure lasted for approximately 2.5 to 3 hours, including the time required to set up the EEG equipment.

### 4.3 Behavioral Analyses

Missing responses (no response within 2000 ms after probe presentation) and incorrect response assignments were considered as errors. Responses faster than 150 ms (i.e., premature responses) after probe onset were excluded from the analyses.

Error rates and RT patterns were analyzed by means of analyses of variance (ANOVA) with the within-subject factors *‘cued items’* (one item vs. two items cued) and *‘non-cued items’* (one item vs. two items non-cued). Only the selective retro-cue conditions were considered for these analyses. Additionally, we compared error rates and RTs between selective and neutral retro-cue conditions by means of within-subject *t*-tests. These analyses only included trials with two-item memory arrays (as neutral retro-cues were only presented in this experimental condition).

### 4.4 EEG recording and preprocessing

The EEG was recorded using 65 Ag/AgCl passive electrodes (Easycap GmbH, Herrsching, Germany) in extended 10/20 scalp configuration with a 1000 Hz sampling rate. A 250 Hz low-pass filter was applied during recording with a NeurOne Tesla AC-amplifier (Bittium Biosignal Ltd, Kuopio, Finnland). Channel FCz served as online reference and channel AFz was used as the ground electrode. Impedances were kept below 10kΩ.

Further analyses were performed using MATLAB and the EEGLAB toolbox^30^. The data were down-sampled to 500 Hz and re-referenced to the average of all channels. Low-pass filters (30 Hz, 33.75 Hz cutoff, 0 to −6 dB transition window) and high-pass filters (0.1 Hz, 0.075 Hz cutoff, 0 to −6 dB transition window) were applied. When kurtosis of a channel exceeded 5 SD (M=6 channels, SD=1.6), it was replaced with a spherical spline interpolation of the immediately proximal channels. Channel rejection and interpolation was not applied to the anterior lateral channels (F9, F10, AF7, AF8, AF3, AF4, Fp1, Fp2), which allowed us to reliably identify eye-movements based on the EEG data. Before creating epochs from 1000 ms before to 3600 ms after memory array presentation, data were again high-pass filtered (1 Hz, 0.5 Hz cutoff, 0 to −6 dB transition window) for later independent component analysis (ICA^54^). Every second epoch was selected and a statistical trial-rejection procedure was used (threshold: 500 µV, probability threshold: 5 SD, max. % of trials to reject per iteration: 5) before running ICA. The resulting ICs were classified by means of ADJUST^55^ and IC weights were transferred back to the whole dataset (0.1 Hz high-pass filter, 30 Hz low-pass filter, average reference, rejected channels). Single dipoles based on a 4-shell spherical head model were calculated for each IC using the DIPFIT plugin for EEGLAB. ICs were excluded from the signal when residual variance of the dipole solution exceeded 50%. Additionally, ICs labeled as vertical eye movements, eye blinks or generic data discontinuities by means of ADJUST were excluded. These analyses were followed by a statistical trial rejection procedure (threshold: 500 µV, probability threshold: 5 SD, max. % of trials to reject per iteration: 5). Finally, trials with strong EEG correlates of horizontal eye movements (saccades) were excluded. This was done based on the signal from lateral frontal electrodes F9 and F10 (i.e., in-cap positions closely matching the hEOG electrode positions on the outer canthus of each eye). A time window of 100 ms was moved in 10 ms steps across each trial, starting at memory array onset and ending at the onset of the memory probes (i.e., 2400 ms later). A trial was rejected, when the voltage difference at F9 or F10 between the first and the second half of at least one of the 100 ms time windows was greater than 20 µV. This led to an additional rejection of 0 to 194 trials (M=74, SD=62.839) per dataset^56^.

### 4.5 Time-frequency analysis

Event-related spectral perturbations (ERSP^30^) were computed by convolving 3-cycle complex Morlet wavelets with each epoch of the EEG data. Resulting time-frequency values consisted of 200 time points between −582 and 3182 ms referred to memory array onset and frequencies ranging from 4 Hz to 30 Hz in 52 logarithmic steps. The pre-stimulus interval served as spectral baseline. The number of cycles in the data window increased half as fast as the number of cycles used in the corresponding fast-fourier transformation (FFT). This led to 3-cycle wavelets at lowest frequency (i.e. 4 Hz) and 11.25-cycle wavelets at highest frequency (i.e. 30 Hz). The channels rejected during preprocessing were re-computed by means of spherical channel interpolation included in EEGLAB prior to these analyses. The same criteria were used for calculating ERSPs on IC level (see below).

#### 4.5.1. EEG correlates of attentional selection in working memory

Posterior hemispheric asymmetries in alpha power in response to the retro-cues were assessed as oscillatory correlates of the orienting of attention on the level of visual working memory representations. These effects were measured by calculating the oscillatory power in the alpha frequency range (8 – 14 Hz) separately for electrodes contralateral vs. ipsilateral to the position of cued vs. non-cued memory items presented on the lateral positions (PO7/PO8, PO3/PO4, P7/P8, P5/P6; see ^45,57^). In a first step, these contralateral vs. ipsilateral responses were assessed in the data collapsed across all selective retro-cue conditions (i.e., one item cued / one item non-cued; one item cued / two items non-cued; two items cued / one item non-cued; two items cued / two items non-cued). Then the contralateral minus ipsilateral difference in alpha power elicited by cued items (with non-cued items on central positions) was compared to the hemispheric asymmetry elicited by non-cued lateral items (with cued items on central positions). Within 1000 permutations, these experimental conditions (cued vs. non-cued items lateral) were randomly assigned for each dataset and two-sided *t*-tests were calculated for all 200 ERSP time points. This resulted in a 200 (time points) by 1000 (permutations) matrix. The largest cluster of *p*-values < 0.01 was measured for each permutation. Differences in hemispheric alpha power asymmetries in the original data were considered as statistically significant when cluster sizes with *p*-values < 0.01 were larger than the 95^th^ percentile of the permutation distribution of maximum cluster sizes. Post-hoc ANOVAs with within-subject factors for *‘eliciting stimulus’* (cued vs. non-cued items lateral), the number of cued items (one vs. two) and the number non-cued items (one vs. two) were run based on the significant clusters. The same cluster permutation approach was applied for comparing the hemispheric alpha power asymmetries elicited by cued and non-cued items with the neutral retro-cue condition.

#### 4.5.2. EEG correlates of motor planning

To achieve a better isolation of the neural activity related to motor planning processes in the left and right sensorimotor cortex, a clustering procedure was applied on IC level. Only ICs with in-brain dipole solution and a residual variance below 20% were included in the cluster analysis. Based on these criteria, each dataset contributed 18.125 ICs on average (range 3 to 29). A k-means clustering algorithm (MATLAB statistics toolbox) separated the overall 435 ICs into 18 clusters and one additional cluster including outlier ICs. These outliers were defined by ICs with more than 3 SD from any of the cluster centroids. This number of clusters was chosen, as it guaranteed a sufficient dissociation of sensorimotor mu/beta suppression from posterior alpha power effects, while also ensuring a high number of datasets contributing ICs to the mu/beta clusters. The clustering was based on IC dipole solutions (3 dimensions), frequency spectra from 4 to 30 Hz (10 dimensions), scalp maps (10 dimensions) and ERSPs from 4 to 30 Hz based on the whole segment (10 dimensions). The number of dimensions define to what extent the different features contribute to the generation of the clusters. As the IC dipole solution can only contribute 3 dimensions (x, y and z values), its relative contribution to clustering was weighted by a factor of 10.

Based on these criteria, we observed two IC clusters that featured the typical characteristics of the mu and beta oscillatory response in preparation for responses and during their execution. There was one left-hemispheric or contralateral mu/beta cluster and one right-hemispheric or ipsilateral cluster. Both clusters featured the typical spectra with a double-peak in higher alpha and beta frequencies (see figure 7). The neural sources of these clusters were estimated by first specifying the MNI coordinates with the peak dipole density and then expanding this point to a sphere with a radius that had the length of 1 SD referred to each dipole coordinate (x, y and z). The statistical sources were defined by the number of grid points within this extended spatial sphere that belonged to a specific anatomical structure, divided by the number of all grid points (see figures 4 and 5).

**Figure 7.**
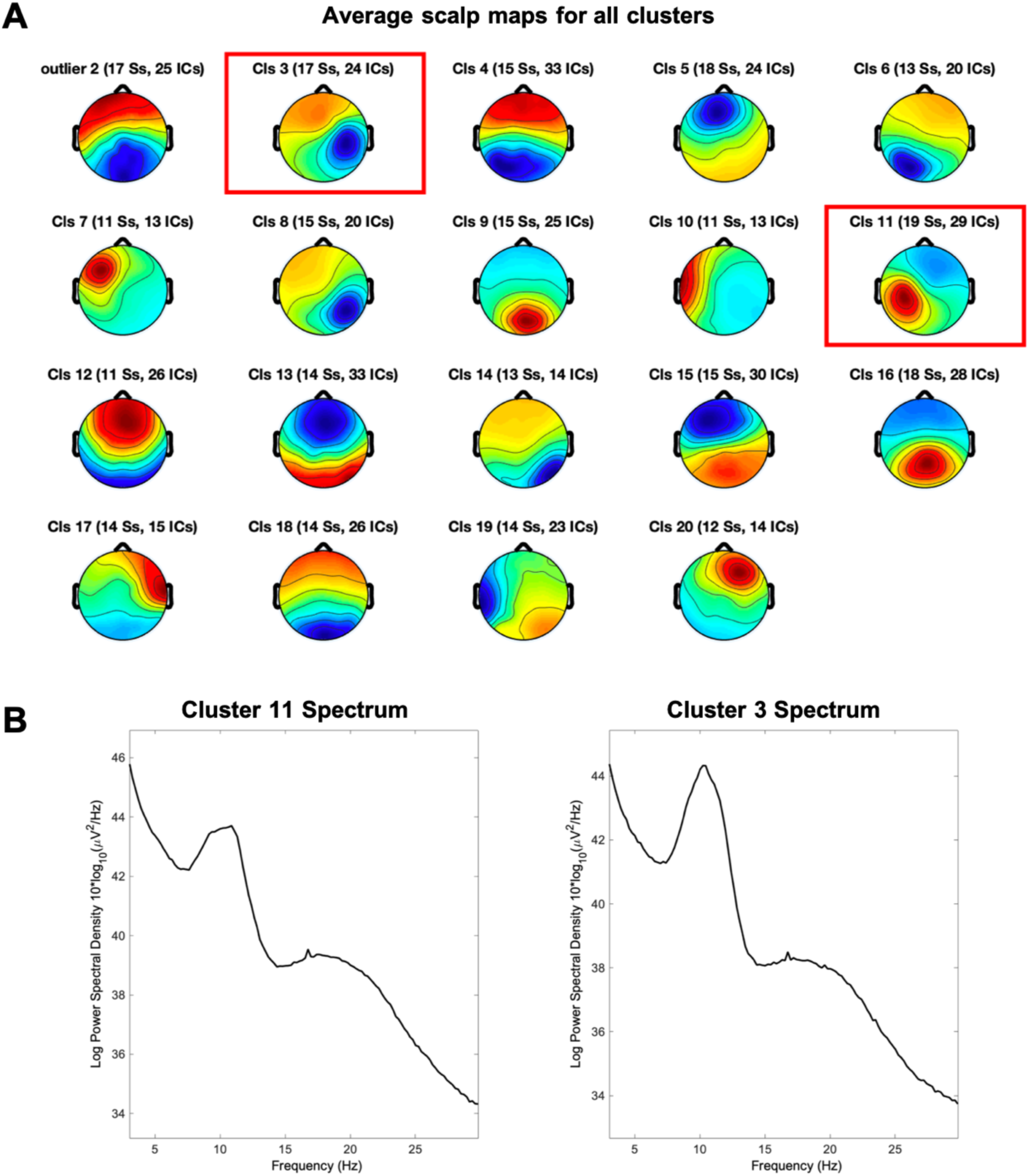
Average scalp maps for all IC clusters (A). The left- and right hemispheric mu/beta clusters are highlighted. 7B depicts the spectral response of these two clusters with a clear peak in the alpha(mu) frequency range and a broader spectral response in beta frequencies.

ERSP responses generated by the statistical sources from 19 participants (left-hemispheric cluster) and 17 participants (right-hemispheric cluster) were compared between experimental conditions with one vs. two cued items and one vs. two non-cued items. A cluster-permutation permutation statistical account was applied to assess time-frequency areas that differed reliably as a function of these experimental conditions. Based on 1000 permutations, the experimental conditions (one vs. two items cued; one vs. two items non-cued) were randomly assigned for each dataset. Afterwards, within-subject *t*-tests were run for each time-frequency data point. The result were two matrices (one for each comparison) with 52 (frequencies) × 200 (times) × 1000 (permutations) *p*-values. The size of the largest cluster of time-frequency points with *p*-values < 0.01 was assessed for each permutation. Significant differences between conditions in the original data were defined as clusters with *p*-values < 0.01 that were larger than the 95^th^ percentile of the permutation-based distribution of maximum cluster sizes (see solid lines in figures 4 and 5).

#### 4.5.3. Brain-behavior correlations

Each ERSP data point within the left-hemispheric mu/beta cluster (averaged across all experimental conditions) was correlated with the average individual RT based on a Spearman rank correlation (i.e., a between-subjects correlational approach). Time frequency clusters featuring reliable correlations between RTs and oscillatory power were assessed with a cluster-permutation approach: First, the assignment of individual RTs to ERSPs was randomized within 1000 permutations. Spearman’s rho was calculated for each time-frequency point within each permutation. Afterwards, we determined for each rho value the probability to belong to the distribution of time-frequency specific rho-values across permutations. Within each permutation, we then measured the sizes of clusters with probability values < 0.01 and stored the maximum cluster size. For the original data, we also assessed the probability of each rho-value to belong to the distribution of time-frequency specific rho-values in the permuted data and searched for clusters with p < 0.01. A cluster in the original data was considered as statistically significant when it was larger than the 95^th^ percentile of the permutation-based distribution of maximum cluster sizes. Afterwards, Spearman rank correlations were again calculated with average RTs and the mean of the time-frequency values within the significant clusters.

## 5. Acknowledgements

We would like to thank Isabel Skiba for contributing to data collection and Tobias Blanke for programming the experiment.

